# *Arfgef1* haploinsufficiency in mice alters neuronal endosome composition and decreases membrane surface postsynaptic GABA_A_ receptors

**DOI:** 10.1101/743625

**Authors:** JiaJie Teoh, Narayan Subramanian, Maria Elena Pero, Francesca Bartolini, Ariadna Amador, Ayla Kanber, Damian Williams, Sabrina Petri, Mu Yang, Andrew S. Allen, Jules Beal, Sheryl R. Haut, Wayne N. Frankel

## Abstract

*ARFGEF1* encodes a guanine exchange factor involved in intracellular vesicle trafficking, and is a candidate gene for childhood genetic epilepsies. To model *ARFGEF1* haploinsufficiency observed in a recent Lennox Gastaut Syndrome patient, we studied a frameshift mutation *(Arfgef1*^fs^) in mice. *Arfgef1*^fs/+^ pups exhibit signs of developmental delay, and *Arfgef1*^fs/+^ adults have a significantly decreased threshold to induced seizures but do not experience spontaneous seizures. Histologically, the *Arfgef1*^fs/+^ brain exhibits a disruption in the apical lining of the dentate gyrus and altered spine morphology of deep layer neurons. In primary hippocampal neuron culture, dendritic surface and synaptic but not total GABA_A_ receptors (GABA_A_R) are reduced in *Arfgef1*^fs/+^ neurons with an accompanying decrease in GABA_A_R-containing recycling endosomes in cell body. *Arfgef1*^fs/+^ neurons also display differences in the relative ratio of Arf6^+^:Rab11^+^:TrfR^+^ recycling endosomes. Although the GABA_A_R-containing early endosomes in *Arfgef1*^fs/+^ neurons are comparable to wildtype, *Arfgef1*^fs/+^ neurons show an increase in GABA_A_R-containing lysosomes in dendrite and cell body. Together, the altered endosome composition and decreased neuronal surface GABA_A_R results suggests a mechanism whereby impaired neuronal inhibition leads to seizure susceptibility.

**Highlights:**

1. *Arfgef1*^fs/+^ mice have lower seizure threshold but no spontaneous seizure.
2. *Arfgef1*^fs/+^ neurons show reduced dendritic surface GABA_A_R.
3. *Arfgef1*^fs/+^ neurons have decreased GABA_A_R-containing recycling endosome accompanied with an increase in GABAAR-containing lysosomes.

## Introduction

Epileptic encephalopathy (EE) is a group of brain disorders often with childhood onset and accompanied by serious neurocognitive consequences (Jain et al., 2013). While linkage and association studies have suggested that *ARFGEF1* is involved in genetic epilepsy (Wallace et al., 1996; Piro et al., 2011; Addis et al., 2018), it was one of several candidate genes and no specific causal *ARFGEF1* variants were indicted previously.

*ARFGEF1* encodes Brefeldin A (BFA) inhibited guanine-nucleotide-exchange protein-1, also known as BIG1 (Addis et al., 2018); and is highly conserved across mammals and eukaryotes (Wright et al., 2014). Arfgef1 belongs to the GBF/BIG family based on the relatively large molecular size compared to other ARF-GEFs and a distinctive feature – sensitivity to Brefeldin A (BFA) (Yamaji et al., 2000). Among multiple domains, the BFA-sensitive Sec7 catalytic domain is the most studied (Le et al., 2013; Lin et al., 2013; Zhou et al., 2013). Arfgef1 selectively activates Class I ARFs to initiate conversion of ARF-GDP to ARF-GTP during the process of intracellular vesicle formation and trafficking (Zhao et al., 2002). Earlier studies also showed that the N-terminal DCB domain interacts with Arl1, targeting Arfgef1 to the trans-Golgi network (TGN) membrane (Galindo et al., 2016); the C-terminus interacts with kinesin and myosin (Saeki et al., 2005; Shen et al., 2008) as well as regulating cell surface localization of ABCA1 or GABA_A_R (Lin et al., 2013; Li et al., 2014). The role of Arfgef1 in the TGN has been studied in detail (Manolea et al., 2008; Boal and Stephens, 2010; Lowery et al., 2013), where it activates Arf1-GTP to recruit clathrin coats, AP1 and Arf binding proteins, also known as GGAs (Lowery et al., 2013). These vesicle-forming components with cargo-sorting capability form new vesicles and traffic cargo from TGN to endosomes or plasma membrane (Shen et al., 2006).

An earlier study suggested that BIG2 (Arfgef2) is functionally associated with endosomal integrity whereas Arfgef1 is associated with the Golgi (Shen et al., 2006). However, newer evidence suggests that Arfgef1 is also localized on endosomes (D’Souza et al., 2014). Since the main Arfgef1 target, Arf1, localizes on recycling endosomes and regulates trafficking from recycling endosome to plasma membrane (Nakai et al., 2013) and GGAs are known to facilitate trafficking between TGN and lysosomes (Hirst et al., 2000), it is reasonable to suggest that Arfgef1 may have a role in post-TGN endosomal trafficking, similar to that of BIG2. However, this aspect of Arfgef1 is understudied.

Arfgef1 regulates neurite development *in vitro* (Zhou et al., 2013). Recently, different groups demonstrated that Arfgef1 is required both for the survival of cortical deep layer neurons and for the initiation of neuronal myelination (Teoh et al., 2017; Miyamoto et al., 2018). Another *in vitro* study showed that decreased Arfgef1 abundance is coupled with decrease of plasma membrane GABA_A_R without affecting total GABA_A_R, resulting in impaired chloride ion influx (Li et al., 2014). However, it is not yet known directly whether Arfgef1 has a role in phenotypes that rely on maintaining inhibition:excitation balance *in vivo.*

Understanding the underlying disease etiology and mechanisms may provide insight for new therapies, especially needed for severe childhood epilepsies. Here, we examine the effect of *Arfgef1* haploinsufficiency in a new mouse line based on a Lennox-Gastaut patient that carries a *de novo* loss of function mutation in *ARFGEF1.*

## Results

### *ARFGEF1 de novo* nonsense mutation in a Lennox-Gastaut Syndrome patient

*ARFGEF1* was detected as a candidate gene in a boy diagnosed with Lennox-Gastaut Syndrome. Seizures began at 6 months of age, and were subsequently diagnosed as infantile spasms. By 3 years of age, the patient exhibited generalized convulsions as well as myoclonic, tonic, and atonic seizures. Neurological exams showed global developmental delay and axial hypotonia. He had no language and no meaningful vocalizations. He was unable to pull to stand or support his own weight. General physical exam results were normal and no dysmorphic features was observed. The electroencephalogram (EEG) initially showed hypsarrhythmia and infantile spasms. Subsequent EEGs showed the emergence of generalized slow spike-and-wave activity along with paroxysmal fast activity and persistence of multifocal epileptiform discharges, superimposed on a high amplitude, disorganized background. He was treated initially with ACTH, followed by multiple medications including vigabatrin, levetiracetam, clonazepam, zonisamide, phenobarbital, lamotrigine, valproic acid, and rufinamide. He was also treated with the ketogenic diet and underwent a corpus callosotomy. Despite these interventions the seizures remained severely refractory.

Exome sequencing performed as part of the Epi4k Collaborative (Epi4K Consortium et al., 2013) revealed a single nucleotide variant resulting in a premature translational stop codon in exon 30 of the *ARFGEF1* gene (ENST00000262215.3:c.4365C>A; p.Cys1455Ter). The patient also has a missense mutation in the *BMP2* gene (ENST00000378827.4:c.328C>A; p.Arg110Ser), classified as “probably damaging” based on a PolyPhen-2 score of 0.968. However, since *BMP2* variants have not yet been associated with neurological disease in human or mouse, and because *Arfgef1* null mouse mutants have abnormal brain development (Teoh et al., 2017), we considered *ARFGEF1* the more likely candidate.

### Genetic analysis of *Arfgef1* haploinsufficiency

We generated an *Arfgef1* knockout mouse line on the C57BL/6NJ strain background by using CRISPR/Cas9 to target the orthologous region on mouse Chromosome 1 (Fig. 1A; Methods). Line *Arfgef1*^em3Frk^ (shown hereafter as *Arfgef1*^fs^) has a 4-nt frameshift mutation in exon 30, resulting in a premature termination codon (MQYVMYSPSI*) closely mimicking the patient’s mutation. Although heterozygous mice are viable and fertile, all homozygotes died within one day of birth, similar to a previous report of the *Arfgef1* knockout mouse (Teoh et al., 2017). The presence of cDNA and protein in whole brain at embryonic day 17.5 was examined using primer pairs flanking N-terminal DCB domain, Sec7 domain, exon 29-31 (mutation in exon 30) and C-terminal HDS4 domain (Fig. S1). The Arfgef1 protein was not detected in *Arfgef1*^fs/fs^ whole brain lysate using Arfgef1 antibodies with N-terminal epitopes (Fig. 1B). In *Arfgef1*^fs/+^ brain, Arfgef1 protein was lower in abundance than wildtype, consistent with haploinsufficiency. The brain size of E17.5 embryos from *Arfgef1*^fs/+^ were comparable to wildtype except that the *Arfgef1*^fs/fs^ forebrain was notably smaller.

**Figure 1.**
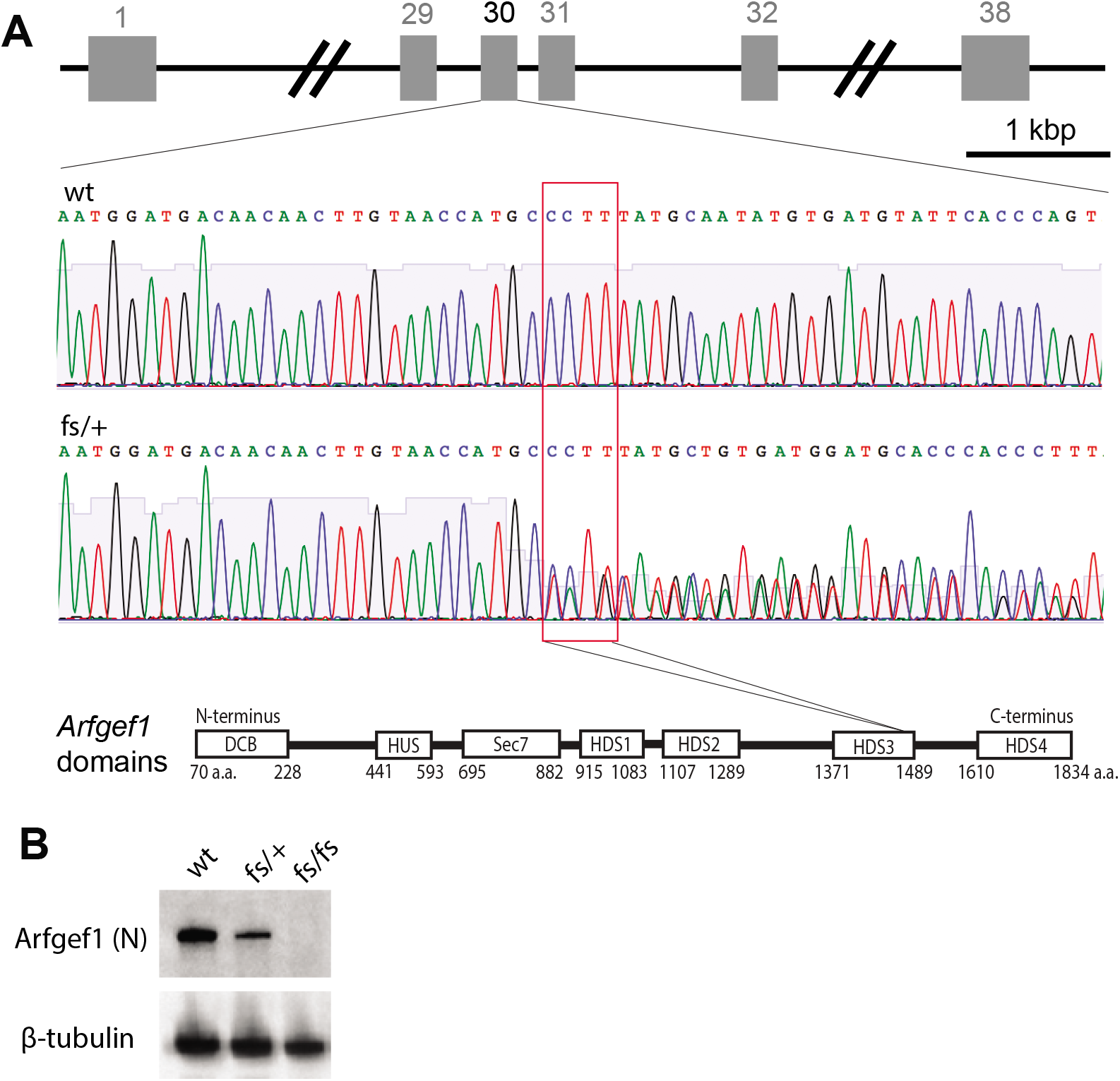
Generation and genotyping of Arfgef1^fs^ mice. ***A***, The CRISPR/Cas9 targeting *Arfgef1* exon 30 on mouse Chromosome 1 resulted in a 4-nt frameshift mutation (*Arfgef1*^fs^). The heterozygotes (*Arfgef1*^fs/+^) were verified with Sanger sequencing. The red box shows the sites of mutation for *Arfgef1*^fs^ correspond to HDS3 domain in Arfgef1 protein. ***B***, The western blot was performed using E17.5 brain lysate (n=3 animals per genotype) and detected using antibody targeting Arfgef1 based on an N-terminal epitope, using ß-tubulin as loading control.

### Developmental delay in *Arfgef1*^fs/+^ mouse pups

We studied the impact of the *Arfgef1*^fs/+^ genotype in mouse pups, to explore parallels with the Lennox-Gastaut Syndrome patient. Newborn pups are known to perform typical behavioral activities that serve as benchmarks for neurodevelopmental delay during postnatal development (Hill et al., 2008). *Arfgef1*^fs/+^ pup body weight did not change significantly from postnatal day (PND) 3 to PND11 (Fig. 2A), suggesting that the feeding behavior was normal. However, in the first week the latency was increased modestly for several tasks, including surface righting reflex and negative geotaxis (Fig. 2B, C), while the vertical screen holding was similar to wildtype (Fig. 2D, E). This is consistant with the hypotonia phenotype found in the patient. Pups younger than PND12 emit ultrasonic vocalization (USV) when separated from dam and littermates (Ferhat et al., 2016). The USV calls in wildtype pups peaked at PND6 and then gradually diminished. In contrast, the peak and diminishment of USV calls was significantly delayed in *Arfgef1*^fs/+^ (Fig. 2F). This indicates developmental delay and is consistent with what was seen in patient. Representative samples of USV calls from wildtype and *Arfgef1*^fs/+^ pups are shown in Fig. S2.

**Figure 2.**
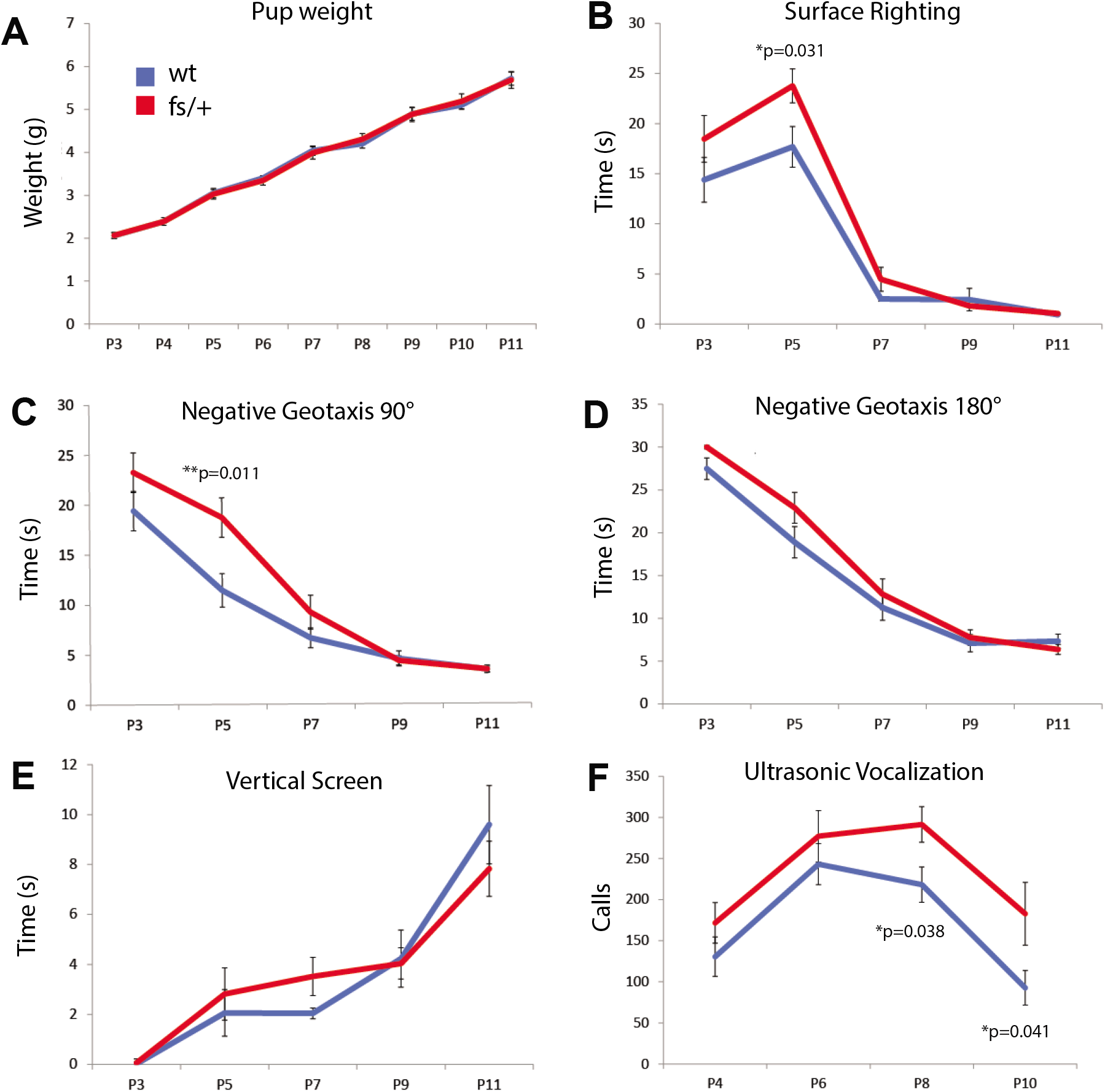
Developmental milestone and ultrasonic vocalization of Arfgef1^fs/+^ pups. ***A***, The weight of *Arfgef1*^fs/+^ and wildtype pups from PND3 to PND11 is statistically similar. ***B, C***, The time required for surface righting, negative geotaxis 90° of *Arfgef1*^fs/+^ pups is statistically slower than wildtype at PND5. ***D, E***, There is no difference observed in the time for *Arfgef1*^fs/+^ and wildtype pups to accomplish negative geotaxis 180° task or to hold on a vertical screen. ***F***, The ultrasonic calls are statistically higher at PND8 and PND10 for *Arfgef1*^fs/+^ pups compared to wildtype pups. Data are collected from pups of both sexes. wildtype, n=22 animals; *Arfgef1*^fs/+^, n=23 animals from 7 litters. Permutation *p*-values shown were determined using the Wilcoxon rank-sum test.

### Morphological defects in *Arfgef1*^fs/+^ brains and neurons

An earlier study showed that the *Arfgef1* knockout heterozygotes have normal brain size during embryogenesis (Teoh et al., 2017), but the postnatal brain was not examined. First, we observed that *Arfgef1*^fs/+^ mice showed transient weight loss by PND14, and catch-up to wildtype by PND45 (Fig. 3A). Using this as a guide for histological examination, we then compared *Arfgef1*^fs/+^ and wildtype brain at PND14, PND30 and PND45 (Fig. 3B). The overall brain size of *Arfgef1*^fs/+^ was similar to wildtype, but dysregulated growth in the dentate gyrus lining of *Arfgef1*^fs/+^ hippocampus was apparent from PND30 (Fig. 3B, red arrowheads). We then examined the morphology of single neuron by diolistic labeling using DiI, a lipophilic dye that diffuses through the membrane lipid bilayer and stains the whole cell structure, first in primary neuron culture and then in brain slices. At 14 days *in vitro* (DIV), we observed long, filamentous and tertiary branch-like structures with larger growth cones on the dendrite of *Arfgef1*^fs/+^ primary hippocampal neurons (Fig. 3C, red arrowheads). These tertiary structures did not adhere to the culture dish surface compared to the respective primary dendrite. Larger growth cone size has been shown to correlate with slower neurite growth rate (Ren and Suter, 2016), suggesting a delay in dendritic branch maturation. In *Arfgef1*^fs/+^ brain slices, although the total spine density was similar in deep layer neurons of both genotypes (Fig. 3D, 3E), the distribution of spine morphological subtypes was altered. In *Arfgef1*^fs/+^ dendrites, the fraction of mushroom-shaped spines was significantly higher than observed in wildtype (Fig. 3D, bottom inset, red arrowheads), whereas there were fewer thin spines and filopodia (Fig. 3F). Since mushroom spines on cortical deep layer neurons are typically more persistent than the immature thin spines or filopodia (Trachtenberg et al., 2002), this evidence suggested changes in dendritic spine plasticity.

**Figure 3.**
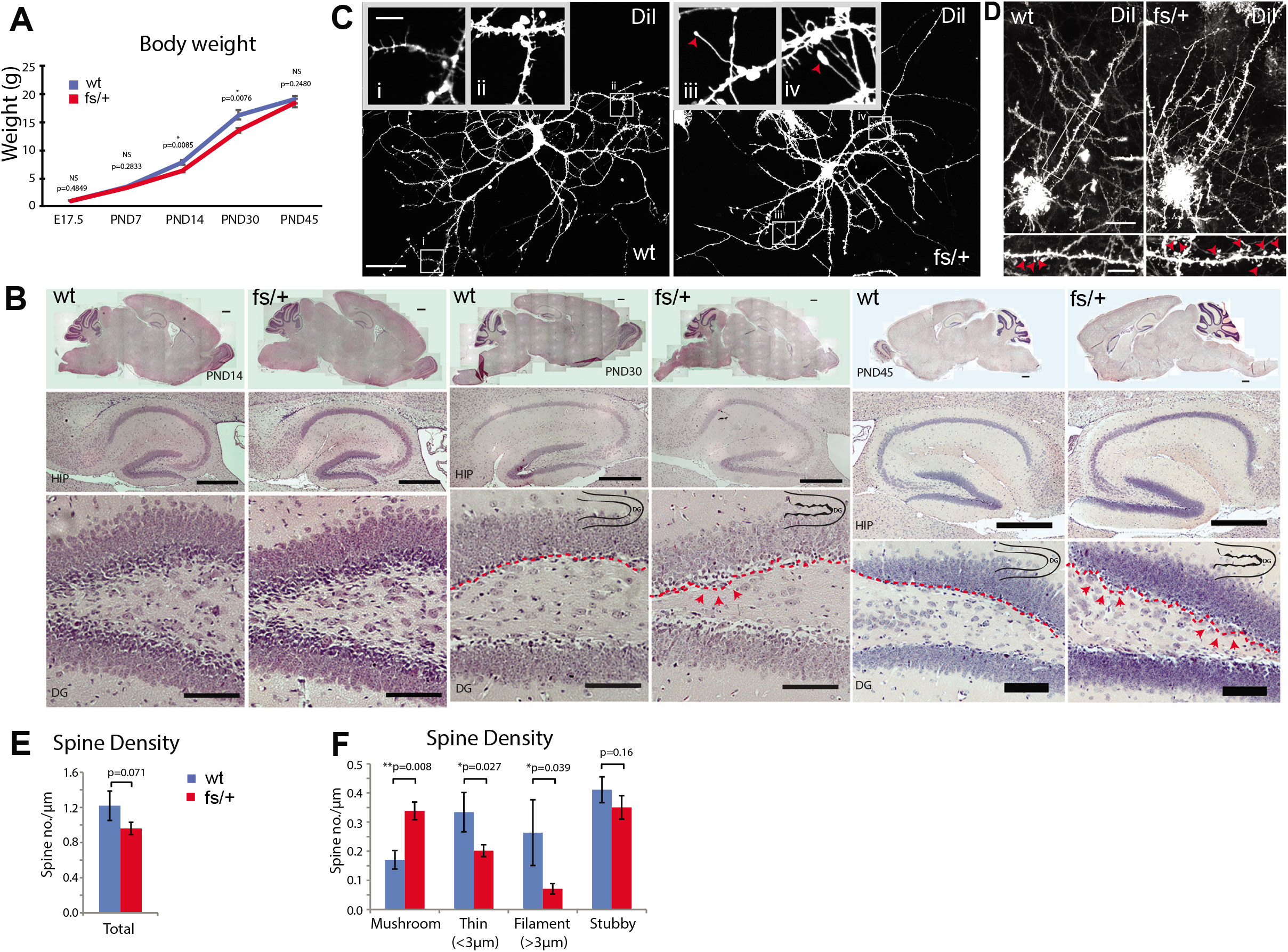
Morphological characterization of Arfgef1^fs/+^ mice. ***A***, The body weight of *Arfgef1*^fs/+^ mice is statistically lower at PND14 and PND30 compared to wildtype mice. E17.5, wildtype, n=16; *Arfgef1*^fs/+^, n=12 from 4 litters; PND14, 30 and 45, wildtype, n=7 mice; *Arfgef1*^fs/+^, n=8 mice from 5 litters. B, The HE-stained sagittal sections shows irregular bumps (red arrowheads) along *Arfgef1*^fs/+^ dentate gyrus lining (red dotted lines) at PND30 and PND45. n=3 animals per genotype for each time-point. Scale bars, sagittal sections, 500 μm; HIP, 500 μm; DG, 100 μm. C, In DIV14 culture, there are filamentous tertiary structures with large growth cones (red arrowheads) appeared on *Arfgef1*^fs/+^ neurons. wildtype, 4 neurons from 1 animals; *Arfgef1*^fs/+^, n=11 neurons from 3 animals. Scale bars, 50 μm; inset, 10 μm. D, Single deep layer neuron is labeled using diolistic method in PND14 deep layer neurons. Mushroom spines were marked by red arrowheads. *E* & *F*, The total spine density is similar between *Arfgef1*^fs/+^ neurons and wildtype neurons. However, spine subtype density (mushroom, thin and filament) are significantly different. Wildtype and *Arfgef1*^fs/+^, n=3 animals each. Scale bars, 20 μm; enlarged box, 10 μm. Data are collected from mice of both sexes. HIP, hippocampus; DG, dentate gyrus.

### Seizure susceptibility of *Arfgef1*^fs/+^ adult mice

In four years of breeding, *Arfgef1*^fs/+^ mice show no signs of spontaneous convulsive seizure throughout their lifetime, nor was there evidence of seizure or epileptiform activity in 48 hr video-EEG recordings of heterozygous adult mice (Fig. S3). However, both male and female heterozygotes have a significantly reduced electroconvulsive threshold to the minimal forebrain clonic seizure endpoint, compared with wildtype littermates (Wilcoxon test with permutation *p*=0.003 and *p*=0.001, respectively; Fig. 4A). They are also susceptible to tonic-clonic seizures induced by subcutaneous GABA_A_R antagonist pentylenetetrazol (PTZ) at a dose (40 mg/kg) near the threshold for wildtype C57BL/6NJ mice. Thus, 92.7% of *Arfgef1*^fs/+^ heterozygotes reached the tonic-clonic seizure or worse endpoint, in contrast to 18.2% of wildtype mice (p<0.0001, Fisher’s Exact Test). The average latency to the first tonic-clonic seizure was also significantly shorter in *Arfgef1*^fs/+^ mice, by an average of 788 sec (Wilcoxon test with permutation *p*<0.001; Fig. 4B).

**Figure 4.**
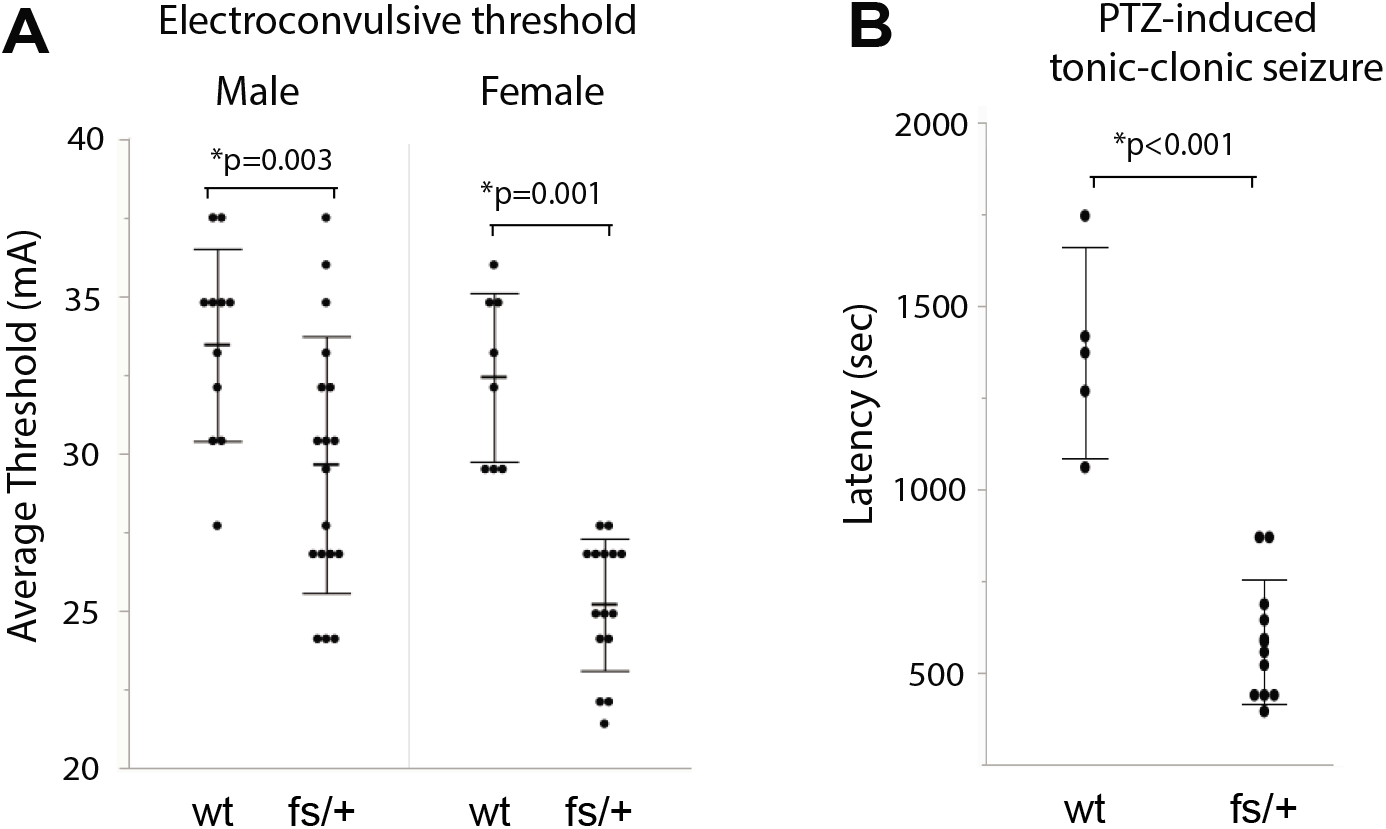
Seizure susceptibility of Arfgef1^fs/+^ mice. ***A***, Electroconvulsive threshold to the minimal clonic seizure endpoint is significantly lower in *Arfgef1*^fs/+^ mice of both sexes (male, wildtype, n=11; *Arfgef1*^fs/+^, n= 18; female, wildtype, n=11; *Arfgef1*^fs/+^, n= 15). ***B***, The latency for PTZ-induced seizure is significantly lower for *Arfgef1*^fs/+^ mice at 9-to 10-week old. wildtype, n=4; *Arfgef1*^fs/+^, n= 11. Permutation *p*-values shown were determined using the Wilcoxon rank-sum test.

### Decreased dendritic surface GABA_A_R in *Arfgef1*^fs/+^ neurons

Li and colleagues (2014) previously demonstrated that siRNA-mediated depletion of Arfgef1 could reduce functional GABA_A_R at the membrane surface (Li et al., 2014). Given the morphological features and seizure susceptibility of *Arfgef1*^fs/+^ mice, we examined GABA_A_R localization in primary hippocampal neuron culture. Through immunofluorescence staining of permeabilized neurons, we determined that the total number of GABA_A_R puncta in proximal dendrite of *Arfgef1*^fs/+^ was similar to that of wildtype (Fig. 5A, D). However, on non-permeabilized neurons, the dendritic surface bound GABA_A_R puncta count decreased in *Arfgef1*^fs/+^ (Fig. 5B, E). GABA_A_R puncta colocalized with synaptophysin, a synaptic marker, also decreased (Fig. 5C, F), together suggesting a decrease in synaptic GABA_A_R density along the dendritic surface. Initial assessment of synaptic properties of hippocampal pyramidal neurons from these cultures at DIV14 showed that *Arfgef1*^fs/+^ neurons had a nominal increase in mean mEPSC frequency and no difference in mIPSC frequency or in mEPSC or mIPSC amplitude (Figure S4A, B, C).

**Figure 5.**
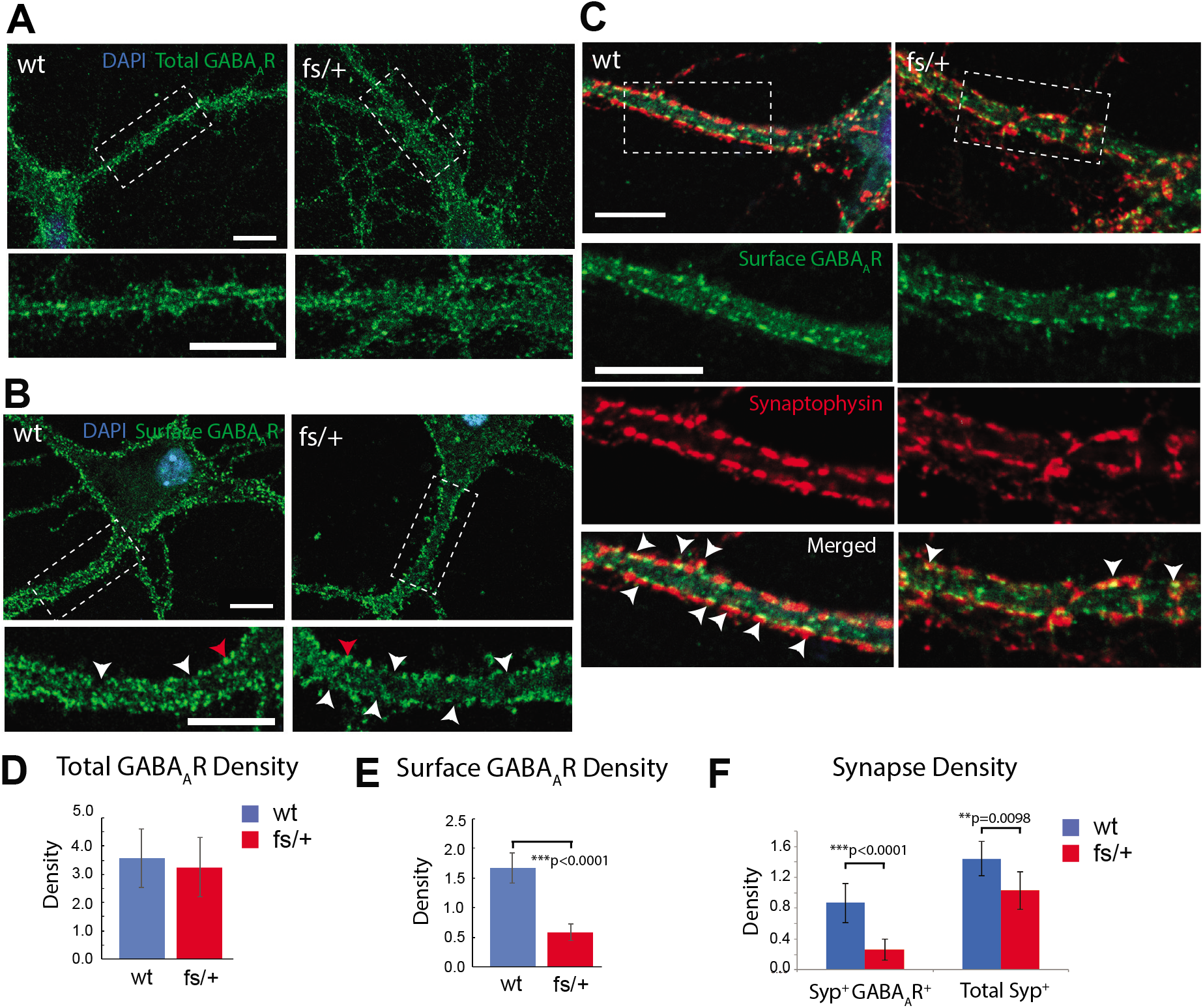
Membrane surface GABA_A_R and synaptic GABA_A_R distribution in Arfgef1^fs/+^ neurons. ***A***, There is no difference in the localization of total GABA_A_R puncta in proximal dendrite. ***B***, GABA_A_R localize on dendritic surface (representative punctum pointed by red arrowheads) at DIV14. The region without GABA_A_R puncta on dendritic surface are marked by white arrowheads Surface GABA_A_R, wildtype, n=19:4 (# neurons:# mice); *Arfgef1*^fs/+^, n=23:7. Total GABA_A_R, wildtype, n=19:4; *Arfgef1*^fs/+^, n=23:7. All scale bars, 10 μm. ***C***, At DIV14, the synaptic GABA_A_R puncta (white arrowheads) on dendritic surface reduced on *Arfgef1*^fs/+^ dendrite. wildtype, n=19:4; *Arfgef1*^fs/+^, n=12:4. All scale bars, 10 μm. ***D, E***, Graphs show quantification of the dendritic surface GABA_A_R density and total GABA_A_R density in dendrite. ***F***, Graph shows the synaptic GABA_A_R density and total synapse density on dendritic surface. Density is defined as punctum number per dendritic length in μm.

### Disrupted distribution of GABA_A_R-containing trafficking vesicles in *Arfgef1*^fs/+^ neurons

Because the total GABA_A_R puncta count remained unchanged, we reasoned that the decrease in synaptic GABA_A_R may result from malfunction either in receptor trafficking or in recycling to the plasma membrane. That is, GABA_A_R might be endocytosed but cannot be recycled back to plasma membrane (Li et al., 2014), or there is slowdown in trafficking of GABA_A_R from Golgi to plasma membrane.

There is significant heterogeneity in the type and subcellular localization of recycling endosomes in neurons, and this heterogeneity is in part represented by Arf6^+^, Rab11^+^, TrfR^+^ markers. For example, Arf6^+^ but not Rab11^+^ and TrfR^+^ recycling endosomes respond to nerve growth factor (NGF) stimulation, suggesting functional differences in recycling endosome species (Kobayashi and Fukuda, 2013). By immunofluorescent staining of primary hippocampal neurons, we determined the density of Arf6^+^, Rab11^+^ and Trf^+^ recycling endosomes by software-assisted segmentation (Fig. S5A, B, C) and quantified vesicle-GABA_A_R colocalization using object-based colocalization (Fig. S7A, B, C, D; see Methods). We found that the the density of Arf6+, Rab11^+^ and Trf^+^ was each decreased in *Arfgef1*^fs/+^ neuronal cell body (Fig. S6Ai, Bi, Ci), as was the density of these respective vesicles that colocalized with GABA_A_R (Fig. 6Ai, Bi, Ci, S7A, B, C). These results suggest either an increase in the dendritic GABAAR recycling, leading to reduction in retrograde trafficking into the cell body, or increase in the recycling endosome that mature into late endosomes. The former is less likely because our result demonstrated reduction in dendritic surface-bound GABA_A_R. Findings in dendrites further reinforced this idea: in *Arfgef1*^fs/+^ dendrites, Arf6^+^ recycling endosomes decreased in density compared to wildtype (Fig. S5A, S6Aii), including those that colocalized with GABA_A_R (Fig. 6Aii, S7A). In contrast, the density of Rab11^+^ and TrfR^+^ recycling endosomes (Fig. S5B, C, 6Bii, Cii) and those colocalized with GABA_A_R (Fig. 6Bii, Cii, S7B, C) in *Arfgef1*^fs/+^ dendrite were comparable to wildtype. These results suggest that *Arfgef1* haploinsufficiency does not only reduced density of Arf6^+^, Rab11^+^, TrfR^+^ recycling endosomes in cell body, such haploinsufficiency may also specifically affects Arf6-dependent dendritic GABA_A_R recycling.

**Figure 6.**
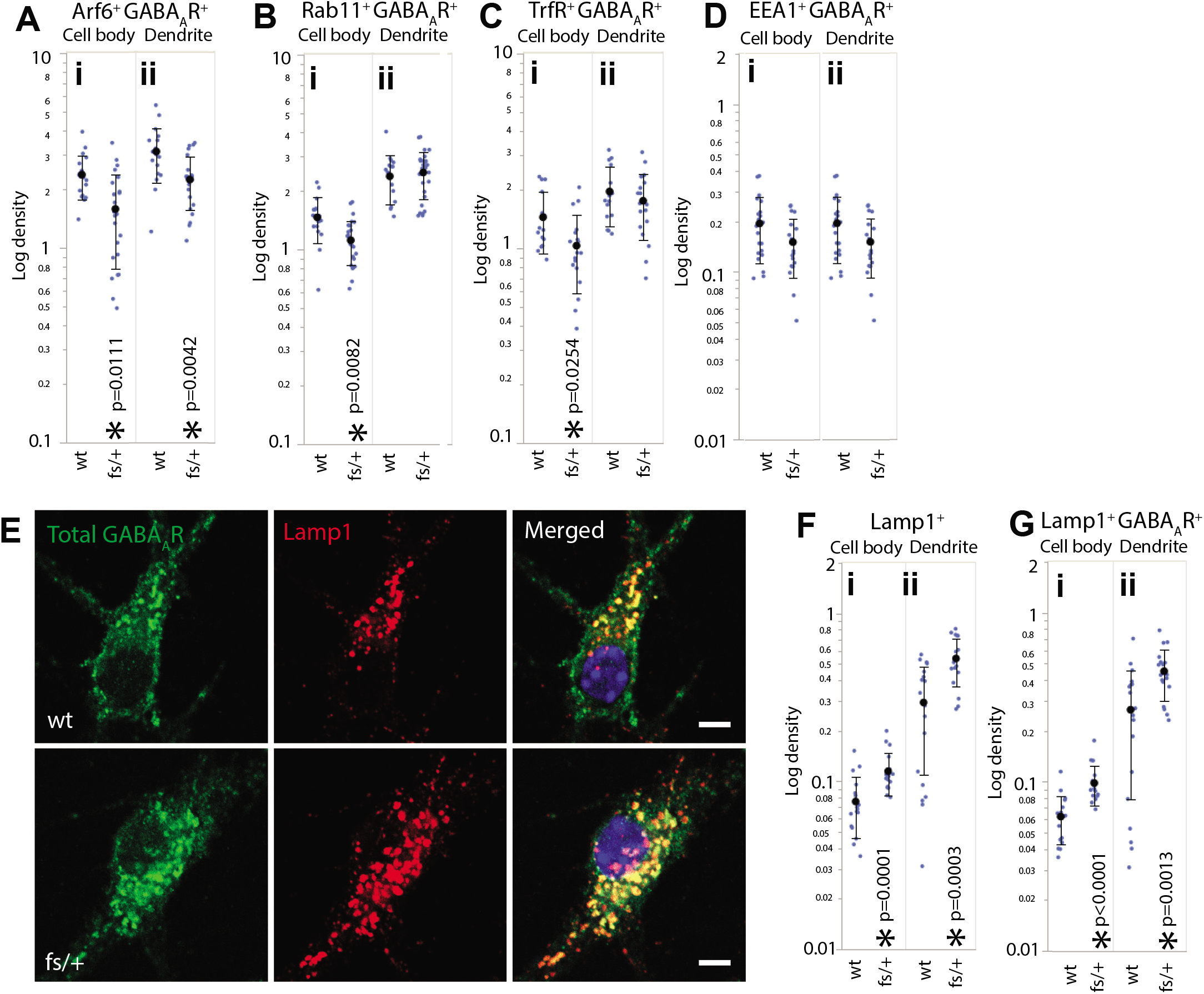
Distribution of intracellular vesicles colocalized with GABA_A_R in wildtype and Arfgef1^fs/+^ DIV14 neurons. ***A, B, C, D***, Graphs show the density of GABA_A_R colocalized with Arf6^+^, Rab11^+^, TrfR^+^ recycling endosomes and EEA1^+^ early endosomes in cell body and dendrite. Densities are displayed in log scale. Density in cell body and dendrite is defined as colocalized punctum number per cell body area (μm^2^) and the punctum number per dendritic length (μm) respectively. Arf6^+^, wildtype, n=19:4 (# neurons:# mice); *Arfgef1*^fs/+^, n=23:7. Rab11^+^, wildtype, n=16:4; *Arfgef1*^fs/+^, n=27:7; TrfR^+^, wildtype, n=15:4; *Arfgef1*^fs/+^, n=18:3; EEA1^+^ wildtype, n=21:6; *Arfgef1*^fs/+^, n=19:5. ***E***, The number of Lamp+ lysosomes and GABA_A_R-containing lysosomes increase in *Arfgef1*^fs/+^ neurons. ***F & G***, Graphs show the density of Lamp1+ lysosome and Lamp1^+^GABA_A_R^+^ lysosomes in the cell body or dendrite. wildtype, n=18:6; *Arfgef1*^fs/+^, n=18:5. All scale bars, 5 μm. Density is shown in log scale. Vesicle densities in cell body and dendrite are defined as punctum number per cell body area (μm^2^) and the punctum number per dendritic length (μm) respectively.

It is useful to consider the impact of *Arfgef1* genotype on the relative ratio of Arf6^+^, Rab11^+^, and TrfR^+^ recycling endosomes, because of the interelationship between these microorganelles. Pairwise comparisons of mutant to wildtype showed that the recycling endosome ratio was significantly skewed in *Arfgef1*^fs/+^ dendrite (p=0.006) but not in cell body (p=0.252). The imbalance in recycling endosome populations in dendrite could be the reason for the decrease of dendritic surface GABA_A_R.

The early endosome is an intermediate organelle, which can either turn into recycling endosomes or mature into lysosomes (Naslavsky and Caplan, 2018). Earlier evidence showed that Arfgef1 also localizes on early endosomes and functions similarly as Arfgef1 localized in the trans-Golgi network (D’Souza et al., 2014). Reduction in the density of early endosomes in dendrite could be the reason for the change in the density of recycling endosomes. However, when compared to wildtype, there were no changes in the density of EEA1^+^ early endosomes in dendrite or cell body of *Arfgef1*^fs/+^ neurons (Fig. S5D, S6Di, ii), as well as the density of early endosomes that colocalized with GABA_A_R (Fig.6Di, ii, S7D).

### Accumulation of GABA_A_R in lysosomes in *Arfgef1*^fs/+^ neurons

The decrease of surface GABA_A_R accompanied by decrease in recycling endosomes suggest that the internalized receptors might have been targeted elsewhere. It has been shown that internalized GABA_A_R that fail to recycle back to the plasma membrane are directed to lysosomes for degradation (Kittler et al., 2004). This led us to determine whether reduction in surface GABA_A_R was accompanied by lysosomal accumulation of GABA_A_R. Indeed, in cell body and dendrite of *Arfgef1*^fs/+^ neurons, Lamp1^+^ lysosome density (Fig. 6E, Fi,ii), as well as lysosome that colocalized with GABA_A_R (Fig. 6Gi, ii) increased compared to wildtype. This showed that GABA_A_R in *Arfgef1*^fs/+^ neurons were redirected to lysosomes instead of being recycled back plasma membrane.

## Discussion

*Arfgef1*^*fs*/+^ mice were generated to model a Lennox-Gastaut Syndrome patient heterozygous for a *de novo* mutation that creates a premature translational stop in the Arfgef1 C-terminus, and thus a haploinsufficient state via presumed nonsense-mediated decay. *Arfgef1*^*fs*/+^ mice do not have spontaneous convulsions, as is sometimes the case with genotypically accurate mouse models of childhood epilepsy that either do not have spontaneous seizures (Frankel et al., 2009; Amendola et al., 2014) or have seizures of a different type than described in the respective patients (Warner et al., 2016; Kovacevic et al., 2018). However, *Arfgef1*^*fs*/+^ do have a low threshold to induced seizures, indicating clear seizure susceptibility. Their abnormal performance in developmental milestone tasks, modest but significant, also suggests global developmental delay and possibly hypotonia, like the patient. The predicted maturation rate for mouse is 150 times faster than human in the first month. Thus, the age at which the *Arfgef1*^*fs*/+^ developmental features are most apparent, between ages PND5 and PND7, is equivalent to a 2-3 year child (Flurkey et al., 2007). This very rapid postnatal maturation in mouse may also partly account for compensatory neurodevelopmental features that prevent a more severe outcome, such as spontaneous seizures.

*Arfgef1*^*fs*/+^ animals have a normal lifespan, whereas *Arfgef1^fs/fs^* die perinatally; in agreement with the earlier report of *Arfgef1* knockout mice (Teoh et al., 2017), except the forebrain defect evident at E17.5 in *Arfgef1^fs/fs^* mice, is more severe. This difference could be due merely to genetic background: the earlier study did not use an inbred strain but rather a mixed 129/SvJ;C57BL/6J background in which hybrid vigor may have mitigated severity.

The GABA_A_R is a membrane-bound ligand-gated ion channel selective for chloride ions (Moroni et al., 2011). Accumulation of GABA_A_R at the synaptic site is required for inhibitory synapse formation and for fast synaptic inhibition (Bogdanov et al., 2006). Other than being trafficked from Golgi to plasma membrane (Luscher et al., 2011), surface GABA_A_ receptors also undergo constitutive endocytosis and then are recycled back to the membrane surface or delivered to lysosome for degradation (Kittler et al., 2000). Our results suggest that both copies of *Arfgef1* are required to maintain normal endosome composition, which is in turn critical for receptor recycling.

Arl1 recruits Arfgef1 to early endosomes (D’Souza et al., 2014). Internalized membrane surface receptors fed into early endosomes have several fates. Surface receptors stay in early endosomes if they are not quickly recycled to plasma membrane from early endosomes or delivered into recycling endosomes for slow recycling. The early endosomes with surface receptors that are not destinied for recycling then mature into lysosomes for degradation (Kittler et al., 2004; Naslavsky and Caplan, 2018). In *Arfgef1*^fs/+^ neurons, there was no change in dendritic early endosomes density but Arf6^+^ recycling endosomes were available albeit decreased in density, suggesting that less early endosomes are destinied for recycling – as supported by our observation of decreased GABA_A_R on the *Arfgef1*^fs/+^dendrite surface – these vesicles mature into lysosomes, which, in turn, are more abundant in the *Arfgef1*^fs/+^ cell body.

The imbalance of recycling and early endosome populations may also explain why there are more lysosomes in *Arfgef1*^fs/+^. AP1 and GGAs, recruited by the Arfgef to the trans-Golgi network, are required to regulate lysosomal enzyme trafficking from the trans-Golgi to lysosomes (Le Borgne and Hoflack, 1998; Hirst et al., 2000; Lowery et al., 2013). Depletion of Arfgef1 alone does not affect the level of GGAs in the trans-Golgi network (Boal and Stephens, 2010; Miyamoto et al., 2018). It has also been demonstrated that the number of synaptic GABA_A_Rs increase in GGA-deficient mouse (Walker et al., 2016). However, depletion of Arfgef1 reduces the formation of AP-1 complex. Notably, such depletion does not entirely eliminate AP-1 (Miyamoto et al., 2018). Since Arfgef2 can compensate for Arfgef1 in the trans-Golgi network (Ramaen et al., 2007), AP-1 and GGAs levels are required for maintaining the network through to lysosomal trafficking, thus the degradation rate would be be maintained in *Arfgef1*^fs/+^ neurons. In our study, although there is significantly less dendritic surface-bound GABA_A_R, some GABA_A_R can still be transported to the membrane surface. Thus, the reduction of surface GABA_A_R is most likely due to reduced recycling, leading to increased GABA_A_R accumulation in lysosomes.

We can only speculate why *Arfgef1* haploinsufficiency in mouse is less severe than the human disease it models, or why more *ARFGEF1* variants have not been found in epilepsy. Lennox-Gastaut Syndrome is a genetically heterogeneous complex childhood epilepsy, if a primary gene is known at all. *Arfgef1* homozygous knockouts are very severely impaired. One possibility is that a very delicate balance of gene dosage, combined with a limited window of late gestational or early postnatal vulnerability, allows the more rapid mouse postnatal development to push through. A more discrete possibility is that *ARFGEF1* is actually a modifier mutation, requiring another predisposing variant for severe disease. In multiple prior studies, *ARFGEF1* was reported as a candidate together with at least one other gene. For example, in a large Australian family spanning three generations in which *CRH* had been proposed as a EE candidate gene (Wallace et al., 1996), Piro and colleagues later suggested that *ARFGEF1* was also plausible (Piro et al., 2011). In another study of copy number variants (CNV), the authors reported an EE patient with onset of 10 years with *ARFGEF1* and *CSPP1* as candidate genes (Addis et al., 2018). In our study, the Lennox-Gastaut patient also carried a nonsynonymous *BMP2* variant; while BMP2 is not yet associated with epilepsy at least one study has shown that BMP2 signaling promotes the differentiation of some GABAergic neurons (Yao et al., 2010). Regardless, the specific impact of *Arfgef1* haploinsufficiency on endosomal balance and dendritic vesicle fate is an attractive pathogenic mechanism and therapeutic target opportunity that may apply to other forms of childhood epilepsy.

## Materials and Methods

### Generation of *Arfgef1* mice

All mice breeding and husbandry were done within the Institute of Comparative Medicine in Columbia University. The mice were kept under a 12-hr day-night cycle with unlimited access to water and food. Mice at two to six-months-old were used for mating. All animal-related procedures were performed according to the guidelines of the Institutional Animal Care and Use Committee (IACUC) of Columbia University. The germline mutation mice were generated in the Genetic Engineering and Technology Core at The Jackson Laboratory (Bar Harbor, ME). In mouse, the *Arfgef1* gene is on Chromosome 1 and consists of 38 exons. Briefly, a guide RNA directing Cas9 endonuclease together with a single-stranded repair template targeting exon 30 was injected into single-cell mouse embryos of C57BL/6NJ line. Founder line were genotyped using PCR (primer F3, 5’-GTCTGAAGTGAAGCACGTTGG-3’ and primer R3, 5’-CAGTGGGGTCAACGTGTTATG-3’), restriction enzyme digestion (MwoI; New England Biolabs, R0573L) and DNA sequencing (primer F2, 5’-GTGGCTAGAGAGGCTCGTTTT-3’). Identified founder mice were crossed with wildtype C57BL/6NJ, resulting *Arfgef1*^fs^ line (4-nt deletion in exon 30).

### RT-PCR

Total RNA was collected from wildtype and *Arfgef1*^fs/+^ brains at embryonic day 17.5 (E17.5) using TRIzol reagent (Ambion, 15596026). Reverse transcription was performed using SuperScript^TM^ III First-Strand Synthesis SuperMix for qRT-PCR (Invitrogen, 11752-050) according to the manufacturer’s protocol. PCR was then performed using primers flanking *Arfgef1* cDNA corresponding to the exons 3-6 (N-terminal DCB domain), exons 13-17 (Sec7 domain), exons 29-31 (mutation site) and exons 31-38 (C-terminal HDS4 domain) with primerd flaking actin as control. The PCR products were subjected to electrophoresis on 2% agarose gels. The primers used in this experiment were listed below.

DCB domain

Forward: 5’-CAC CCT TCC ACC AGT GAA ATC A −3’

Reverse: 5’-TGC TGC CGA TGT CTT TCT CTT TC −3’

Amplicon size: 495bp

Sec7 domain

Forward: 5’-GGA ATT GGC AGC TAC AGT ACA CAG −3’

Reverse: 5’-CCT GTG GAC TGT GAA GGT CTG −3’

Amplicon size: 505bp

Exon 29-31

Forward: 5’-GAC AAC ATG AAA TTG CCA GAA CAG C −3’

Reverse: 5’-GAT ATC CAG GGT GCA GTT ACA CG −3’

Amplicon size: 285bp

HSD4 domain

Forward: 5’-GCT GGA ACT CAT CCA GAC CAT C −3’

Reverse: 5’-TGA GCT TTA AAC CTG CTG TCA CTG −3’

Amplicon size: 524bp

Actin

Forward: 5’-ACG ATA TCG CTG CGC TGG −3’

Reverse: 5’-GAG CAT CGT CGC CCG C −3’

Amplicon size: 72bp

### Antibodies

The following primary antibodies were used: Anti-Arfgef1 N-terminal epitope (ThermoFisher Scientific, PA5-54894; Santa Cruz, sc-376790; sc-50391), anti-Arfgef1 C-terminal epitope (Bethyl, A300-998A), Anti-GABA A Receptor with epitope for extracellular domain of the β2,3 subunits (Millipore Sigma, MAB341), anti-synaptophysin (ThermoFisher Scientific, PA1-1043), anti-lamp1 (DSHB, 1D4B), anti-eea1 (R&D Systems, AF8047), anti-transferrin receptor (Abcam, ab84036), anti-Rab11 (ThermoFisher Scientific, 71-5300), anti-Arf6 (ThermoFisher Scientific, PA1-093), anti-map2 (Millipore Sigma, Ab5622), anti-ankG (Santa Cruz, sc-28561), anti-ß-tubulin (Proteintech, 10094-1-AP and 66240-1-Ig), anti-GAPDH (Proteintech, 60004-1-Ig), anti-lamin B1 (Proteintech, 66095-1-lg). The following secondary antibodies were used: Goat anti-mouse HRP-conjugated IgG (Proteintech, SA00001-1), goat anti-rabbit HRP-conjugated IgG (Proteintech, SA00001-2), Goat anti-mouse Alexa Fluor 488-conjugated IgG (ThermoFisher Scientific, A11017), goat anti-rabbit Alexa Fluor 594-conjugated IgG (ThermoFisher Scientific, A11072), goat anti-rat Alexa Fluor 594-conjugated IgG (ThermoFisher Scientific, A11007), donkey anti-sheep NL557-conjugated IgG (R&D Systems, NL010).

### Western blotting

Brain lysates were collected using the trichloroacetic acid precipitation method. Briefly, the lysates were separated by SDS-PAGE and transferred to Immobilon-PSQ PVDF Membrane (Millipore Sigma, ISEQ00010). The membranes were incubated with primary antibodies overnight at 4°C, followed by incubation with the appropriate HRP-conjugated secondary antibody for 1 hr at room temperature. Samples were washed three times in PBS with 0.05% Tween-20 (PBST) after each step. Signals were developed using Amersham ECL Western Blotting Detection Reagent (GE Healthcare, RPN2106) and visualized using western blot imaging system (Azure Biosystems, Azure C400).

### Video-EEG

Mice of both sexes, aged 5 weeks and 8 weeks were anesthetized through intraperitoneal injection of 400mg/kg 2,2,2-Tribromoethanol (Sigma Aldrich, T48402). Surgery was performed to drill four burr holes. The first two holes located 1 mm anterior to bregma on both sides. The third hole located 2 mm posterior to bregma on the left side. The fourth hole located over the cerebellum as reference. Four teflon-coated silver wires soldered on a microconnector (Mouser electronics, 575-501101) were placed between dura mater and pia mater and a dental cap was applied. Post-operative mice were given 5mg/kg carprofen (Zoetis, Rimadyl injectable) subcutaneously and a recovery time of 48 hours before electroencephalography (EEG) recording. The recovered mice were then connected to Natus Quantum programmable amplifier (Natus Neurology, 014160) for 24 hours over one day-night cycle. The mice activity was video-recorded simultaneously using a night-vision-enabled Sony IPELA EP500 camera. Differential amplification recordings were recorded pair-wise between the three electrodes and a referential electrode, resulted in a montage of 6 channels for each mouse. The EEG data was analyzed using Natus Database v8.5.1.

### Seizure susceptibility tests

Electroconvulsive threshold testing was performed in mice 7-10 weeks of age as described previously with minor modification(Frankel et al., 2001)s (Frankel et al., 2001). Briefly, a drop of a topical anesthetic (0.5% tetracaine in 0.9% NaCl) was placed on each eye of a restrained mouse, and current was applied using silver transcorneal electrodes connected to an electroconvulsive stimulator (Ugo Basile model 7801) using the following parameters: 299 Hz frequency 1.6 ms pulse width, 0.2 s pulse duration, variable current. Mice were tested approximately daily in 0.5 mA increments until the minimal clonic forebrain seizure endpoint was reached. For analysis the mean integrated root mean square (iRMS) current is reported for each genotype-sex group.

Chemical seizure susceptibility was performed using subcutaneous injection of mice between 9 and 10 weeks of age using pentylenetetrazol (PTZ), a GABAA receptor antagonist. Mice were observed for 30 min and incidence of and latency to the first tonic-clonic seizure endpoint was recorded. The PTZ dose of 40 mg/kg was predetermined to be near the threshold for the tonic-clonic seizure endpoint in C57BL/6NJ mice of this age.

The Wilcoxon non-parametric rank-sum test with 1000 permutations was used in R to determine permutation *p*-values for elecroconvulsive threshold and for latency to PTZ-induced tonic-clonic seizure.

### Developmental milestones and ultrasonic vocalization

All pups from the same litter were kept with respective mother throughout the experimental period. They were numbered with tattoo on the soles at PND2 and evaluated for neurological reflexes on PND 3, 5, 7, 9 and 11. Each subject was tested at the same three-hour time window of a day. The pups were weighted each day during the test. For righting reflex test, the pup was placed on its back on a flat and hard surface. The time a pup took to right itself on all four paws was recorded. For negative geotaxis test, the pup was placed with head facing downward on a 20 x 20 cm flat wire mesh screen slanted at 45° angle. The time a pup took to turn around and face the angle of 90° and 180° from the starting downward direction was recorded. For vertical screen grasping test, the pups were placed on a 20 x 20 cm flat wire mesh standing vertically at an angle of 90° to a flat surface. The time a pup was able to maintain itself on the mesh wire was recorded. All the times were recorded using a stopwatch for a maximum of 30 seconds (s). All test were carried out under room temperature. The pups were returned to the mother immediately after the tests.

Ultrasonic vocalization from a pup being isolated from its mother and littermates was recorded on PND 4, 6, 8 and 10. Briefly, a random pup was removed gently from its nest into an isolation container filled with clean bedding. The container was then placed into a sound attenuating chamber equipped with an UltraSoundGate Condenser Microphone CM 16 (Avisoft Bioacoustics, 40011). The microphone connected to an UltraSoundGate 116 USB audio device (Avisoft Bioacoustics) was linked to a computer installed with Avisoft RECORDER v2.97, where the acoustic data was recorded at a sampling rate of 250,000 Hz in 16 bit format. The software was programmed to filtered away acoustic frequency lower than 15 kHz to reduce background noise interference. Each pup was recorded for 3 min, weighted, and then returned to nest with mother immediately. The number of calls emitted by the pups was counted by an independent operator blinded from the genotypes. The Wilcoxon non-parametric rank-sum test with 1000 permutations was used in R to determine permutation *p*-values.

### Hematoxylin and eosin staining

Mice of respective ages were perfused. The brains were carefully extracted and cut along the lambda-bregma points. Pieces of brains were fixed in 4% (w/v) paraformaldehyde (PFA) in 0.1 M phosphate buffer (PB) at pH 7.4 overnight at 4°C. After serial dehydration steps, the brains were embedded in paraffin (Sigma, P3808). The brains were sectioned at 5 μm thickness using Leica RM2125 microtome. Every tenth section were collected on a glass slide (Matsunami Glass, SUMGP14) and subjected to HE staining. Briefly, the slices were rehydrated and stained with hematoxylin then counterstained with eosin. These slices were subsequently dehydrated using ethanol and xylene and mounted with coverslip using Permount (Fisher Chemical, SP15-500). Histological images were acquired using a Nikon Eclispse E800M light microscope and NIS-elements v4.51. Images were merged using automated function in Photoshop CC 2015.

### *Ex vivo* single neuron diolistic labeling and spine analysis

The postnatal day (PND) 14 brains were collected and fixed as described above. These brains were transferred into fresh 2% PFA in 0.1 M PB at pH 7.4 and stored at 4°C for up to two months before use. The brains were embedded in 4% agarose and shot with a gene gun (Biorad, 1652451) loaded with homemade bullets consisting tungsten microbeads (Biorad, 1652267) coated with DiI (Invitrogen, D282) at 80 PSI. After gentle washing, the brains were immersed in fresh 2% PFA in 0.1 M PB at pH 7.4 and stored in a dark moisten chamber for 3 days at room temperature to allow dye diffusion. Finally, the brains were sliced at 300uM using a vibrotome (Leica, VT1200) and mounted with coverslip using glycerol. Z-stacks images were acquired immediately using Zeiss LSM-800 confocal microscope and Zen v2.3. The quantification and morphological analysis of spines were done using NeuronStudio v0.9.92. In agreement with literature, the spine shapes were categorized as filopodia, thin, mushroom and stubby (Qiao et al., 2016). Spines with a neck are classified as thin or mushroom based on the head diameter. Spines longer than 3 mm were classified as filopodia. Spines without a neck are classified as stubby.

### Primary hippocampal neuron culture

Primary hippocampal neurons were collected according to Fath’s method with little modification (Fath et al., 2009). Briefly, hippocampi were dissected from embryonic day (E) 17.5 embryos, dissociated through trituration with the aid of papain (Worthington, LK003178) pre-incubation. A total of 1 x 10^5^ cells was plated in 35 mm culture dish containing acid-treated coverslips coated with poly-D-lysine (Sigma-Aldrich, P7886) and laminin (Gibco, 23017015). The primary hippocampal neurons were kept in culture medium containing Neurobasal-A medium (Gibco, 10888022) supplemented with B27 (Gibco, 17504044) and Glutamax (Gibco, 35050061). Cytosine β-D-arabinofuranoside (Sigma-Aldrich, C1768) was added once to a final concentration of 3 μM on the next day after 50% medium change. The cultures were maintained at 37°C with 5% CO_2_ in a humidified incubator. Every three days, 50% expensed medium were replaced with fresh culture medium.

### Immunofluorescence

DIV14 neurons were fixed for 10 min at 4°C with 4% (w/v) PFA in 0.1 M PB at pH 7.4. The fixed neurons were incubated in blocking buffer containing 5% goat serum (Gibco, 16210064) and 0.1% saponin (Sigma-Aldrich, 47036) in PBS pH 7.4 (Gibco, 10010031) at room temperature for 30 min. Blocked neurons were subsequently incubated with primary antibodies in blocking buffer at 4°C overnight followed by incubation with appropriate secondary antibodies and DAPI in blocking buffer at room temperature for 2 hrs. Samples were washed three times in PBS after each step. The stained neurons on coverslips were mounted on glass slides using Fluoromount-G (Southern Biotech, 0100-01) and sealed with nail polish after drying at room temperature overnight. Z-stacks images were acquired using Zeiss LSM-800 confocal microscope and Zen v2.3. The quantification and analysis of labeled vesicle and GABA_A_R puncta were done using Imaris v9.2.1 software. Due to the limitation of the point spread function (PSF) for tiny objects in confocal microscopy, colocalization is quantified using an object-based method (Dunn et al., 2011). Briefly, vesicle and GABA_A_R puncta stained with antibodies in cell body or dendrite are recognized and segmented in different channels using Imaris v9.2.1. One GABA_A_R colocalized with one vesicle is defined as one GABA_A_R punctum localizes within 1 μm proximity of one vesicle punctum, center to center. There were no significant differences found in the dendrite length and the cell body area where the data were collected. Therefore, the number of puncta in dendrite was normalized using dendrite length (μm) and the number of puncta in cell body was normalized using cell body area (μm^2^). The normalized numbers were presented as density (in dendrite, vesicle number per μm; in cell body, vesicle number per μm^2^).

For statistical analysis of puncta counts, a log-Poisson mixed model was run using the lme4 package in R (www.R-project.org), with cell body area or dendrite length as a respective covariate and a random effects term included to correct for significant overdispersion in the count data. To determine whether endosome ratios differed between mutant genotype and wildtype, genotype x vesicle type interaction terms were included in a similar model which also included a fixed term for the individual biological from which ratios examined. Statistical comparisons in this study were considered significant if the *p*-value is less than 0.05.

### Single cell patch-clamp recordings

To assess synaptic activity, electrophysiological recordings were carried out from dissociated hippocampal neurons plated at a density of 50000 cells on 15 mm diameter glass coverslips and cultured for 13-15 days. Recordings were made in voltage clamp whole-cell configuration using a Multiclamp 700B amplifier and a Digidata 1550 digital-to-analogue converter (both from Molecular Devices) at a 10 kHz sample frequency. Patch pipettes were fabricated with a P-97 pipette puller (Sutter Instruments) using 1.5 mm outer diameter, 1.28 mm inner diameter filamented capillary glass (World Precision Instruments). The external recording solution contained (in mM): NaCl 145, KCl 5, HEPES 10, Glucose 10, CaCl_2_ 2, MgCl_2_ 2, 0.001 tetrodotoxin, pH 7.3 with NaOH, and osmolality adjusted to 325 mOsm using sucrose. A cesium-based pipette solution contained (in mM): cesium methanesulfonate 130, sodium methanesulfonate 10, EGTA 10, CaCl_2_ 1, HEPES 10, TEA-Cl 10, MgATP 5, Na2GTP 0.5, QX-314 5, pH 7.2 with CsOH, adjusted to 290 mOsm with sucrose. Pipettes had final resistance of approximately 5 MΩ when filled with internal solution. Uncompensated series resistance was <15 MΩ. Series resistance and membrane capacitance were electrically compensated to approximately 70%. Miniature excitatory postsynaptic currents (mEPSCs) and miniature inhibitory postsynaptic currents (mIPSCs), were recorded from the same cell by holding the cells at −60 and 0 mV, respectively. Recordings were acquired for a period of 5 minutes. All recordings were carried out at room temperature (21 – 23 °C). Miniature events were detected offline with Clampfit 12.7 (Molecular Devices) using the template matching function and a minimum threshold of 5 pA. Each event was manually inspected to determine inclusion or rejection in analysis. Further analysis was carried out using R (www.R-project.org).

## Acknowledgements

This research is supported by NIH grant R37 NS031348 to WNF, NIH grant U01HG0009610 to JB and NIH/NIA grant RO1AG050658 to FB. We are grateful to Erin Heinzen-Cox, David Goldstein, Daniel Lowenstein and the EPGP and Epi4k project teams for bringing the *ARFGEF1* variant to our attention, to Osasumwen Virginia Aimiuwu for support in RNA reverse transcription and to Ryan Dhindsa for his R script for determining permutation statistics.

