## Supplementary Figures for "*Arfgef1* haploinsufficiency in mice alters neuronal endosome composition and decreases membrane surface postsynaptic GABA_A_ receptors"

### Figure S1

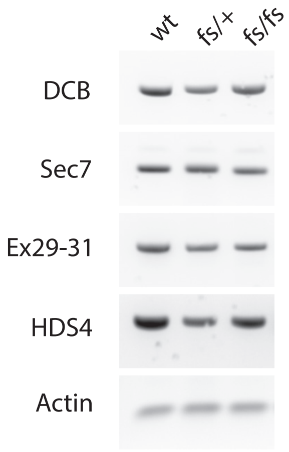


**Figure S1.** *Expression of Arfgef1 RNA.* Representative electrophoresis gel images of PCR products (n=3 animals per genotype). The forward (F) and reverse (R) PCR primers flanking the DCB domain, Sec7 domain, exons 29-31 and HDS4 domain of Arfgef1 cDNA were used for PCR. PCR yielded the expected amplicons as shown.

### Figure S2

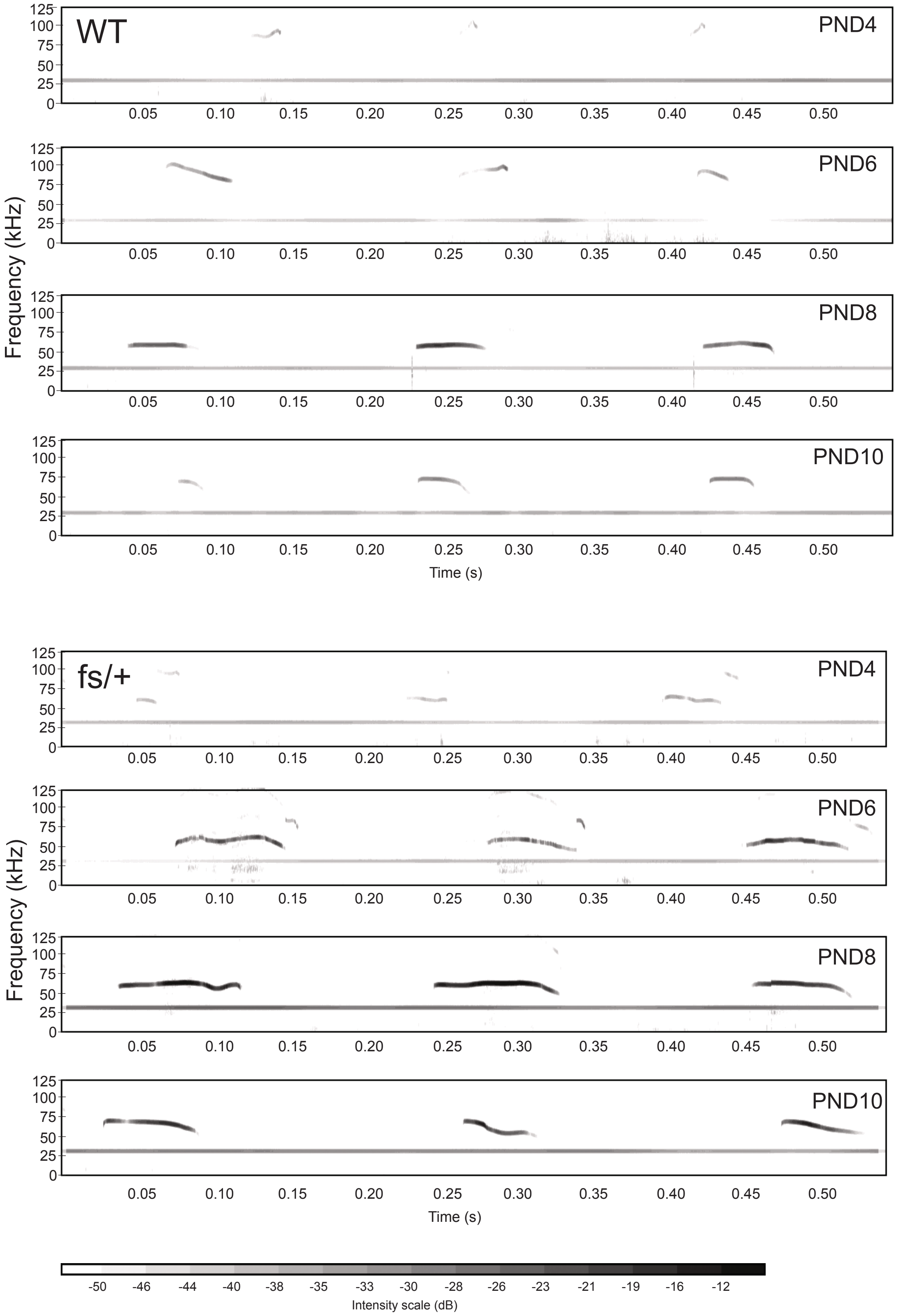


**Figure S2.** *Ultrasonic vocalization (USV) of Arfgef1*$^{fs/+}$ and wildtype pups. Representative USV recordings containing three discrete calls from *Arfgef1*$^{fs/+}$ and wildtype pups at PND4, 8, 10 and 12 are shown. See Fig.2 legend for other details.

### Figure S3

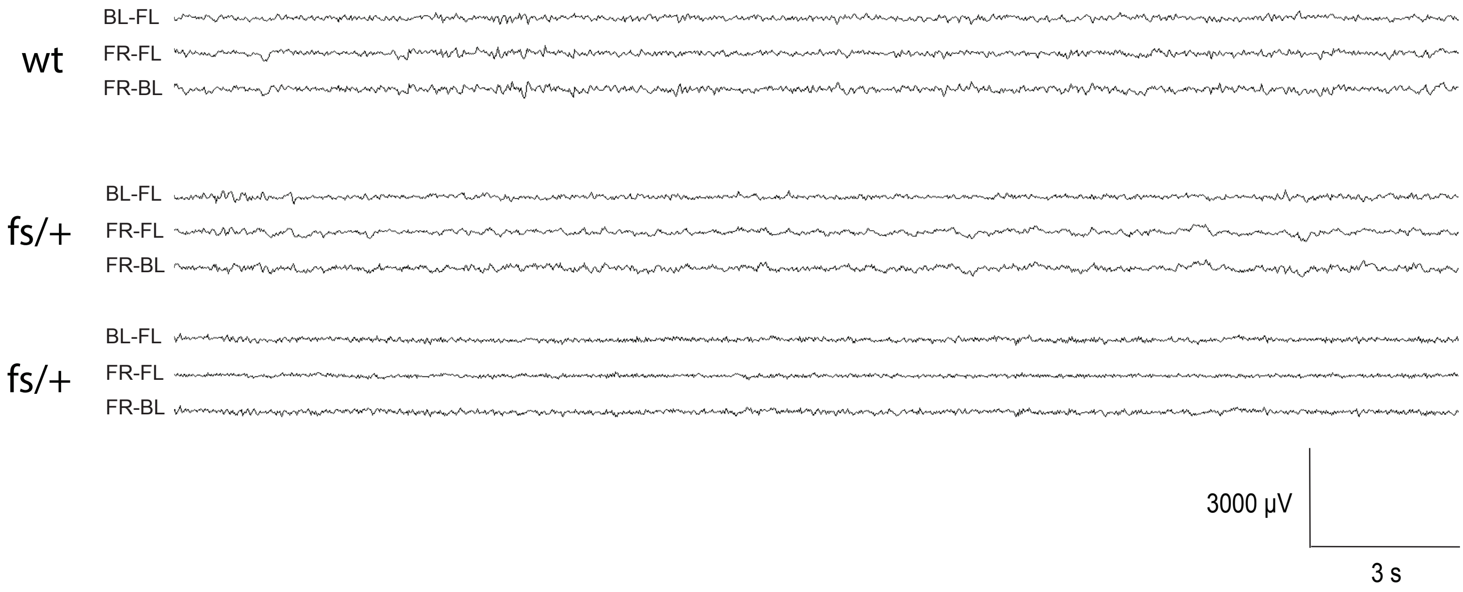


**Figure S3.** *Electroencephalogram (EEG) recording.* Representative EEG recordings at 8 weeks showing no seizure or epileptiform activity in both *Arfgef1*$^{fs/+}$ and wildtype mice. 5-weeks old, wildtype and *Arfgef1*$^{fs/+}$, n=4 animals each; 8-week-old, wildtype and *Arfgef1*$^{fs/+}$, n=3 animals each

### Figure S4

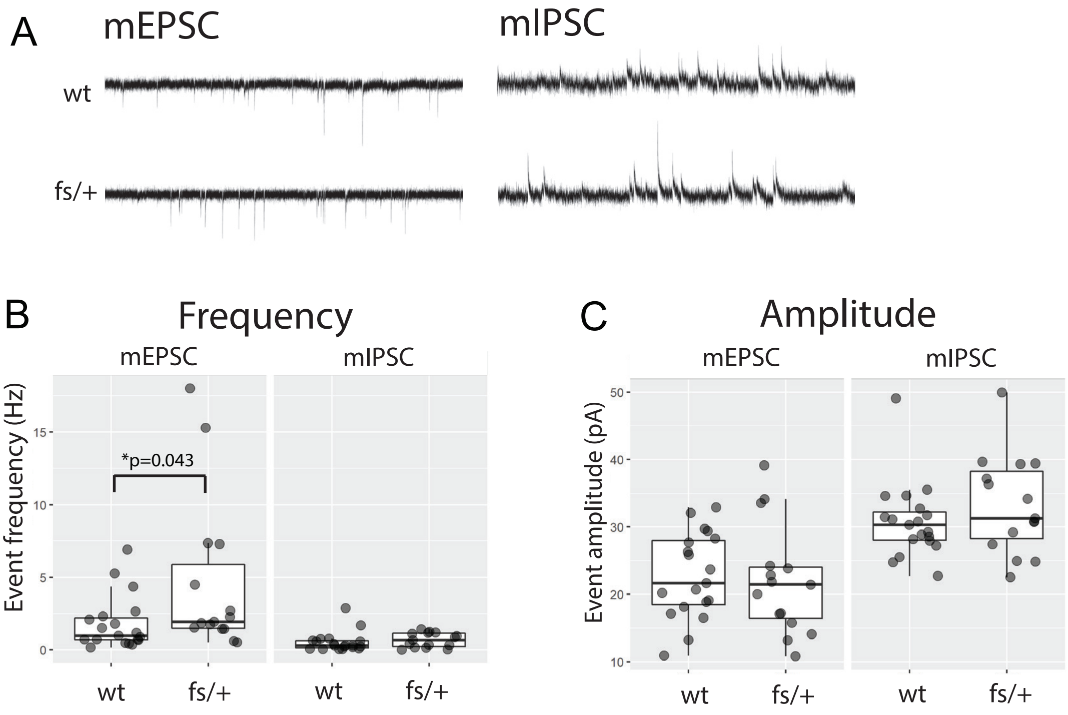


**Figure S4.** *Synaptic activity of Arfgef1*$^{fs/+}$ *hippocampal neurons.* ***A***, Example membrane traces showing mEPSCs and mIPSCs recorded from the same neuron from wildtype and *Arfgef1*$^{fs/+}$ cultures. ***B*** & ***C***, Box plots of frequency and amplitude for mESPC and mIPSC. Each point represents the frequency or amplitude of synaptic currents recording from a single neuron. Box indicates median, first and third quartiles. Whiskers extend to the lowest/highest value within the 1.5 × IQR of hinge. wildtype, n=19 neurons (4 mice); *Arfgef1*$^{fs/+}$, n=15 neurons (4 mice).

### Figure S5

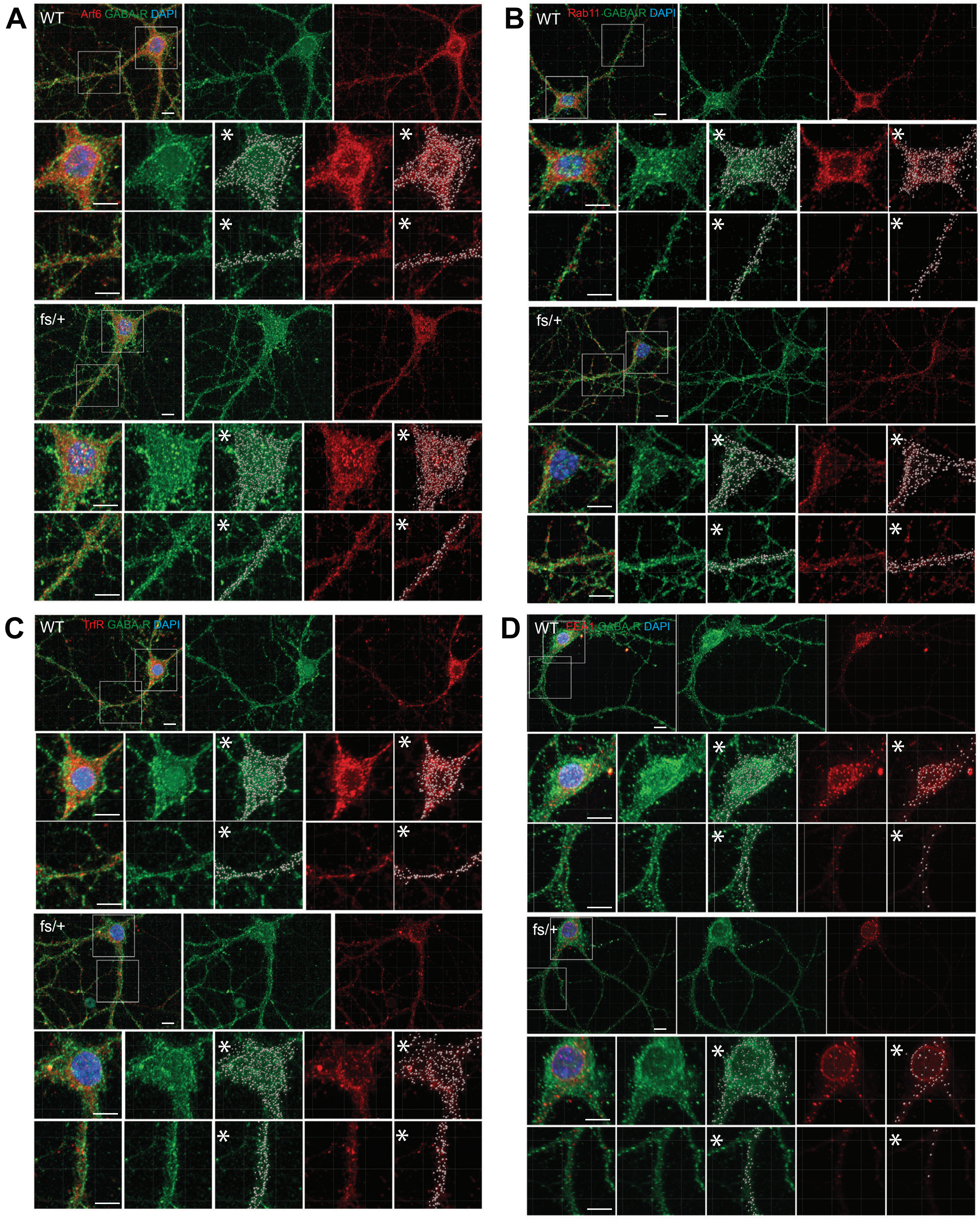


**Figure S5.** *Immunostaining and software segmentation of GABA$_A$ receptor and intracellular vesicles.* ***A***, ***B***, ***C***, ***D***, Maximum intensity projection images generated from confocal z-stack images show the immunostaining of Arf6$^+$, Rab11$^+$, TrfR$^+$ recycling endosome puncta and EEA1$^+$ early endosome puncta in the cell body and dendrite. *Insets show vesicles (red channel) or GABA$_A$R (green channel) in cell bodies and dendrites segmented using Imaris v9.2.1. Each white punctum are counted as one vesicle or one GABA$_A$R. Graphs of the quantification are shown in Fig.S6. Sample sizes are mentioned in Fig. 6.

### Figure S6

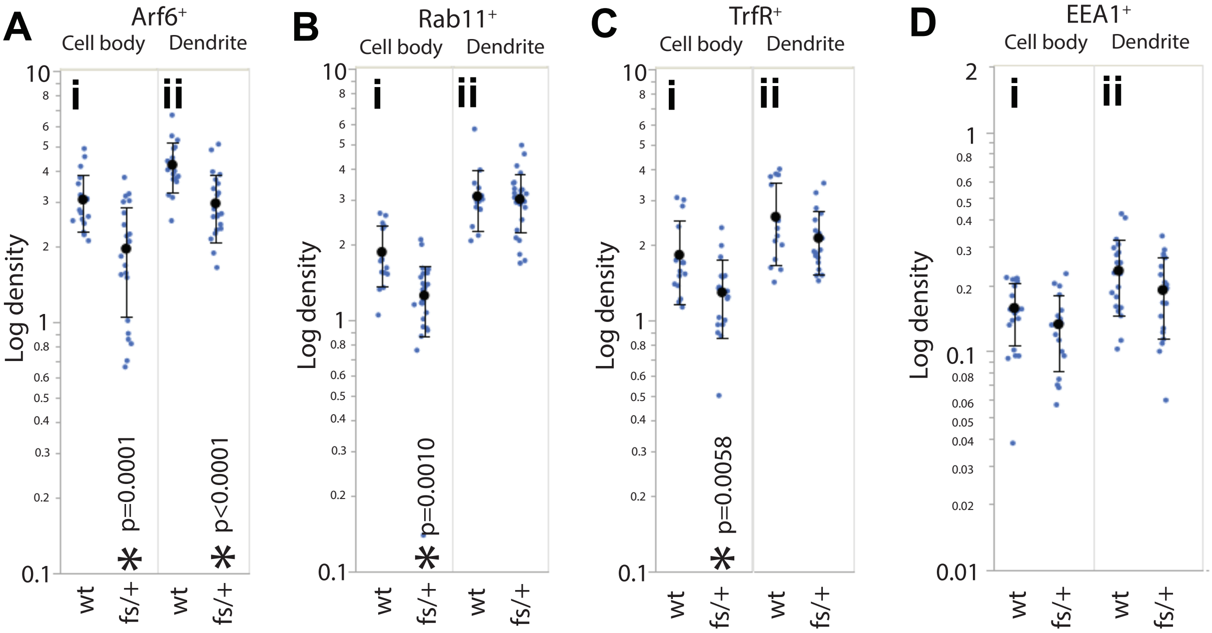


**Figure S6.** *Quantification of intracellular vesicles.* ***A***, ***B***, ***C***, ***D***, Graphs show the density of Arf6$^+$, Rab11$^+$, TrfR$^+$ recycling endosome and EEA1$^+$ early endosome puncta in the cell body or dendrite. Sample sizes are mentioned in Fig. 6.

### Figure S7

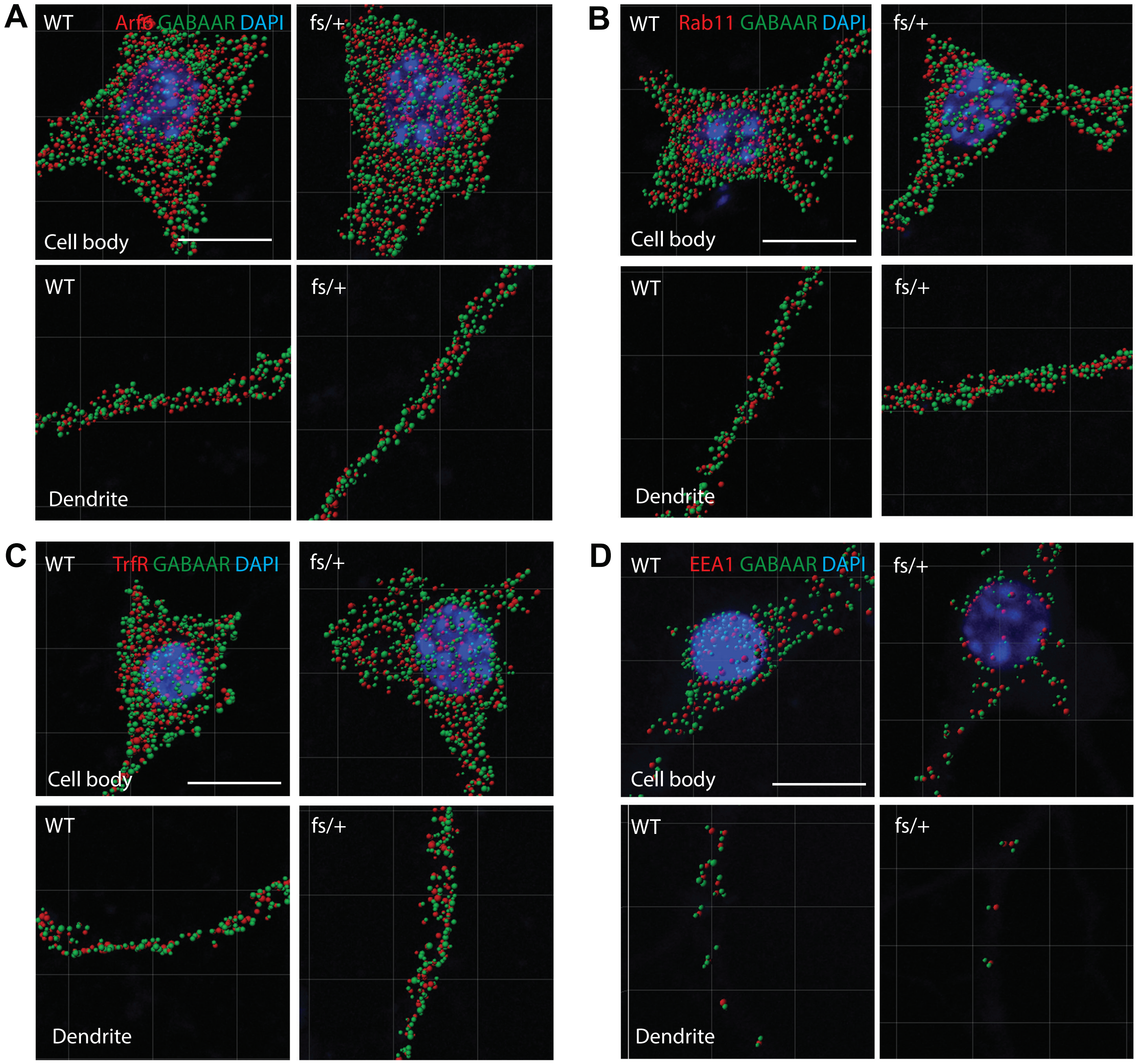


**Figure S7.** *Quantification of object-based colocalization for* $GABA_A$ *receptor and intracellular vesicles.* Representative segmented images of Arf6$^+$, Rab11$^+$, TfrR$^+$ recycling endosome puncta (red puncta) or EEA1$^+$ early endosome puncta (red puncta) colocalized with $GABA_AR$ (green puncta) in the cell body and dendrite. Due to the limited resolution of confocal microscopy, an object-based colocalization method was used for quantification, defined by $GABA_AR$ localizing within 1 µm proximity of a vesicle, center to center.
